# The widespread misconception about the Japanese major biogeographic boundary, the Watase line (Tokara gap), revealed by bibliographic and beta diversity analyses

**DOI:** 10.1101/186775

**Authors:** Shohei Komaki, Takeshi Igawa

## Abstract

The biota of the Japanese Archipelago is divided into the Palearctic and Oriental realms by the Watase line (Tokara gap), a major biogeographic boundary of Japan. This boundary is generally placed between Akusekijima and Kodakarajima Islands of the Tokara Archipelago, and has been the subject of many biogeographic debates. However, despite being widely accepted, the position of the boundary is doubtful because of a lack of clear evidence. Here, to verify the definition and existence of the biogeographic boundary, we performed a documentary search and beta diversity analysis of multiple taxa. Our documentary search suggested that the Watase line (Tokara gap) should be put between Yakushima/Tanegashima and Amamioshima Islands, but recent references to it clearly deviate from its original definition, and that the placement of the boundary line between Akusekijima and Kodakarajima Islands is based on limited and biased evidence. Our beta diversity analyses found no common biogeographic boundary dividing the Tokara Archipelago into two realms, and showed that the beta diversity pattern of this region is explained by the areas and geographic distances of the islands in agreement with the general principles of island biogeography. The widespread misunderstanding of biogeography in this region could have been perpetuated by preconception and the citation of references without verification. Our study proposes that revision of the biogeography in the Tokara Archipelago, a gap region between the Palearctic and Oriental realms, is necessary and demonstrates the negative influence of preconception in biogeographic debate.

## BACKGROUND

The similarity of species components between islands often decreases with the geographic distance between them. This is so-called distance decay, a principle of island biogeography [1]. Island area is also a significant determinant of island biota (the species-area relationship) [2,3]. In the case of land-bridge islands, a geohistory involving sea barrier and land-bridge formation is also an essential factor characterizing the terrestrial and freshwater biota on the island [4-7].

In Japan, there are multiple sea straits whose geohistories seems to have affected the spatial pattern of biodiversity; the Soya, Tsugaru, Tsushima, Tokara and Kerama straits [8-11]. In particular, the Tokara strait is a major biogeographic boundary because it divides the Japanese Archipelago into the Palearctic and Oriental (Indomalaya) realms [10]. This biogeographic boundary is often referred to as the Watase line [12-14]. Alternatively, the term ‘Tokara gap’, which is thought to correspond to the Watase line, is often used in phylogeographic studies [15-18]. In most such studies, this biogeographic or phylogeographic boundary is usually placed between two islands of the Tokara Archipelago, Akusekijima (Akuseki) and Kodakarajima (Kodakara) Islands (gap 5 in Fig. 1) [10,19,20]. It is plausibly explained that migrations of terrestrial and freshwater organisms over the strait have been limited since the Pliocene throughout glacial cycling because of the deep submarine canyon 1000 m below sea level between the two islands [16,20-24] that resulted in the boundary line between the Palearctic and Oriental realms.

**Fig. 1.**
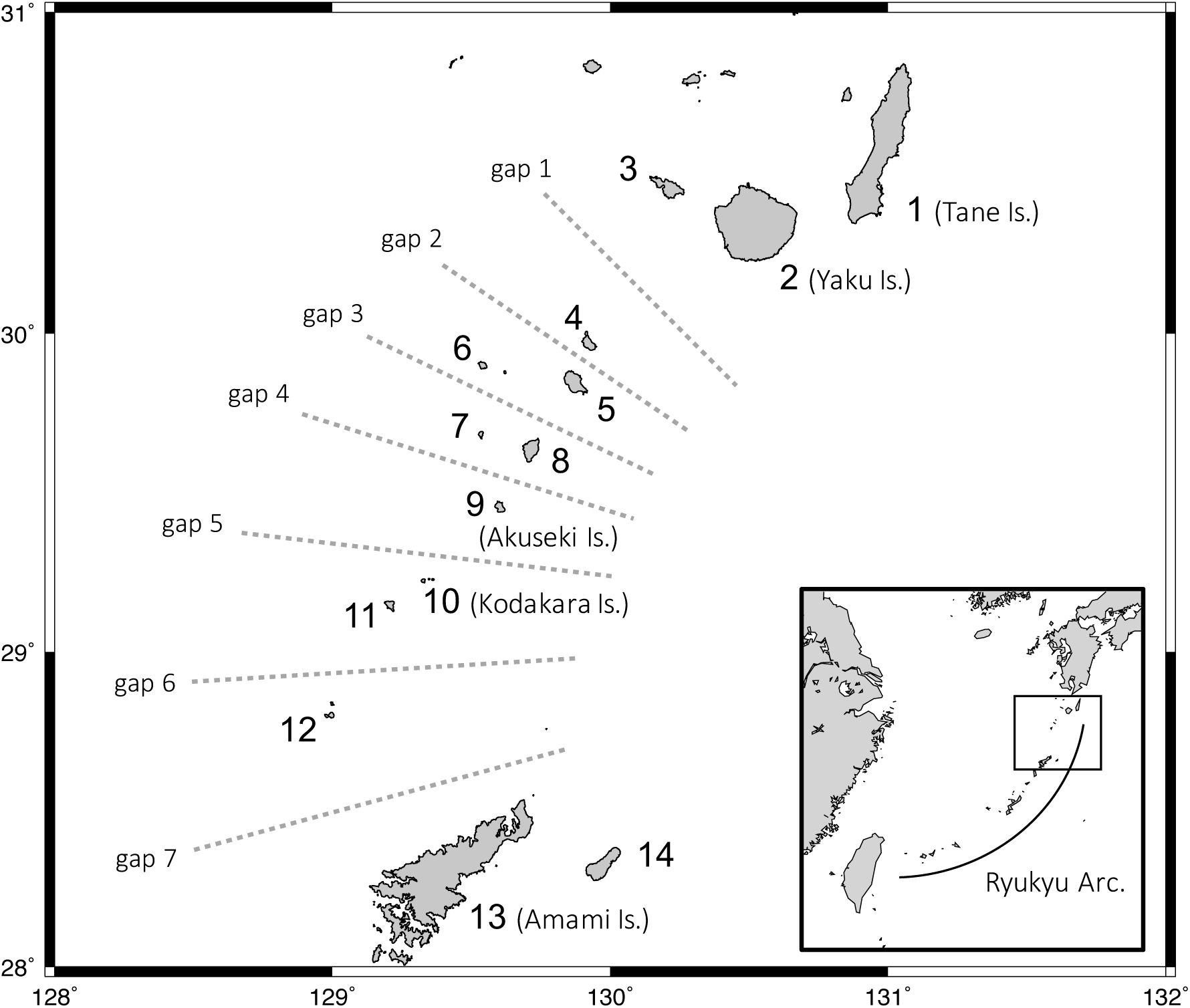
The Tokara Archipelago and adjacent islands. Gaps 1–7 (dashed lines) are hypothetical biogeographic boundaries considered in our analyses. Island numbers correspond to those in Table 1.

Several practical studies of beta diversity have demonstrated that the Tokara gap (gap 5 in Fig. 1) significantly contributed to the biogeographic patterns of terrestrial organisms (Nakamura et al. [20] and Kubota et al. [25] for plant species; Ichikawa et al. [26] for land snails). However, several species distributed across the Tokara gap are considered to have achieved overseas dispersal across the boundary (Kurita and Hikida [15] for skinks; Tominaga et al. [27] for tree frogs). Based on the estimated divergence time between island populations across the Tokara gap, which post-dated the formation of the sea barrier, these studies concluded that the species dispersed over the sea rather than across a land bridge.

Among the abovementioned studies, Ota [9] is the article most frequently referred to in the context of the Tokara gap. Ota [9] reviewed the biogeography of amphibians and reptiles in the Ryukyu Archipelago, and put a boundary line for the Tokara gap between Akuseki and Kodakara Islands in a figure; the idea of this boundary is widely accepted and referred to today. However, why the Tokara gap was put between the two islands was not explained in the article or the references therein; thus, it is unclear whether the boundary line was placed roughly without consideration or on any basis. In addition, a deep submarine canyon is an essential feature characterizing the Tokara gap, but no such canyon exists between Akuseki and Kodakara Islands (Fig. 2). The existence of this biogeographic boundary is, therefore, doubtful despite it being the basis of biogeographic debate in Japan.

**Fig. 2.**
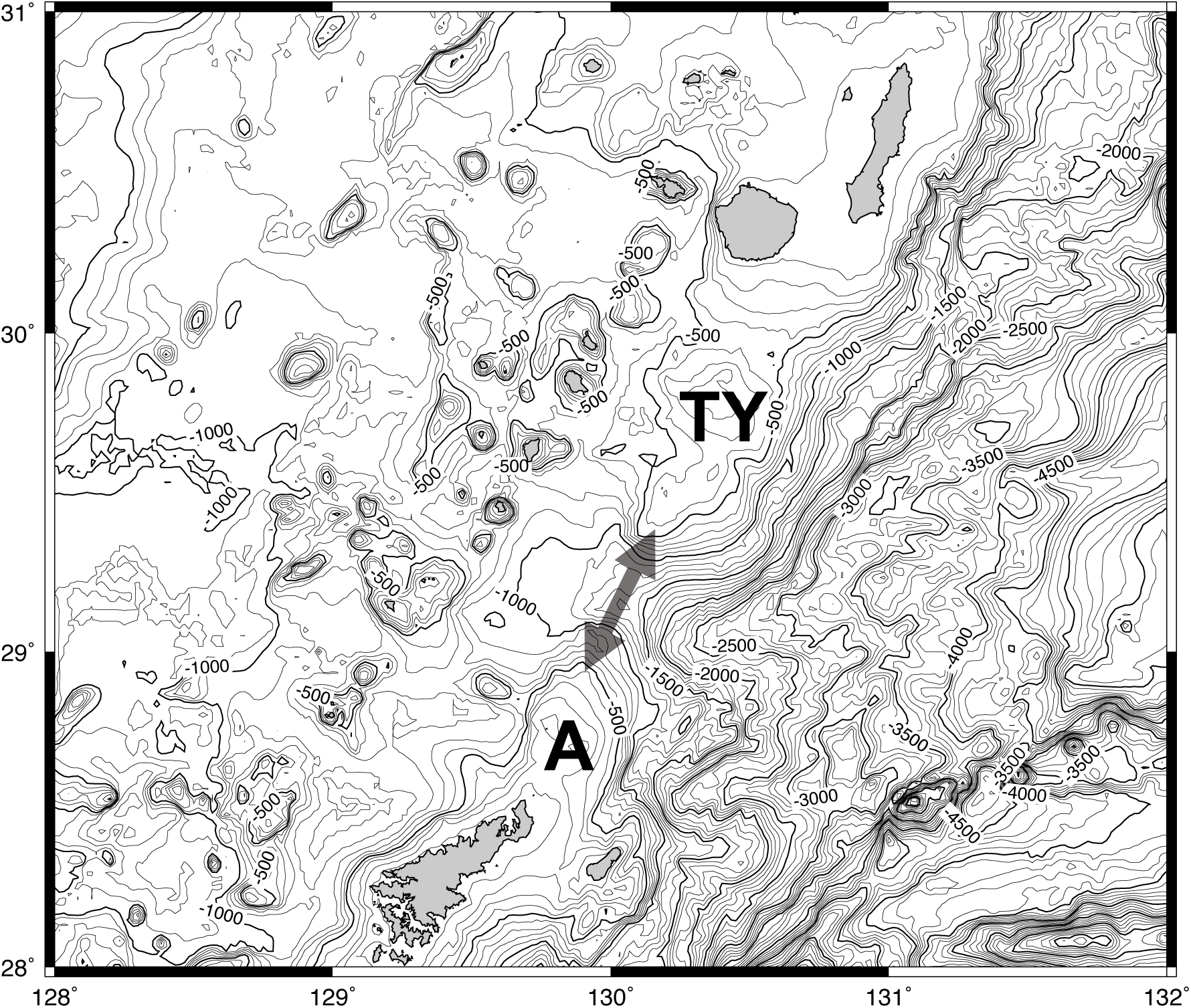
Bathymetric map around the Tokara Archipelago. The arrow represents the Tokara gap, which corresponds to the Watase line. TY: Tane/Yaku Spur, A: Amami Spur.

Here, to understand what the Watase line or Tokara gap is and whether and where it exists, (1) we revisited the concept of these terms in a documentary search, and (2) reanalysed the beta diversities of multiple taxa in the Tokara Archipelago and adjacent islands. Based on the results, we discuss how a biogeographic preconception has spread and affected biogeographic studies in the last decade.

## METHODS

### Document search

Using Google Scholar (https://scholar.google.co.jp), we first searched and read journal articles including original articles, letters, reviews and short notes in which the Watase line (including Watase’s line) or the Tokara gap were mentioned. The Google Scholar search was performed on November 16 2016 using [“Watase line” OR “Tokara gap”] as keywords. Books, proceedings and theses hit by the Google Scholar search were excluded because few documents of this kind are searchable by Google Scholar and thus they do not represent the usage trends of these terms. Instead, documents including articles and books that were referred to as the basis of arguments in any journal articles were checked. Because the term ‘Watase line’ was coined by Dr. Yaichiro Okada after Dr. Shozaburo Watase [13], we also checked related articles published by these authors and references contained therein.

### Biogeographic analyses

Following previous biogeographic studies [20,25,28-30], we used simple and multiple regression models to understand the beta diversity pattern in the Tokara Archipelago. The regression models were based on the dissimilarity of species composition between islands as a dependent variable and hypothetical biogeographic boundaries and environmental factors as explanatory variables. In the analyses, we focused on 14 islands in/around the Tokara Archipelago where the biogeographic boundary is assumed to lie: Tanegashima (Tane), Yakushima (Yaku), Kuchinoerabujima, Kuchinoshima, Nakanoshima, Gajajima, Tairajima, Suwanosejima, Akusekijima (Akuseki), Kodakarajima (Kodakara), Takarajima, Yokoatejima, Amamioshima (Amami) and Kikaijima (Table 1, Fig. 1). The taxonomic groups analysed in this study were land snails [26], ants [29], dragonflies [31], butterflies [32], amphibians [33-35], reptiles [33,36], birds [37-41] and plants [20], as comprehensive and detailed distribution datasets (presence/absence) on each island were available. Uncertain occurrences and artificial introductions were excluded from the datasets. For birds, only the species breeding on each island were considered in the distribution dataset.

**Table 1.** Positions and environmental variables of each island studied.

|  | Island | Latitude (N) | Longitude (E) | Area (km<sup>2</sup>) | Elevation (m) | Forest cover (%) |
| --- | --- | --- | --- | --- | --- | --- |
| 1 | Tanegashima | 30.74361 | 131.0533 | 445 | 282 | 54.4 |
| 2 | Yakushima | 30.33611 | 130.5042 | 504.9 | 1936 | 89.7 |
| 3 | Kuchinoerabujima | 30.44333 | 130.2172 | 35.81 | 657 | 86.3 |
| 4 | Kuchinoshima | 29.96806 | 129.9256 | 13.33 | 628 | 77.1 |
| 5 | Nakanoshima | 29.85917 | 129.8569 | 34.47 | 979 | 77.4 |
| 6 | Gajajima | 29.90306 | 129.5417 | 4.07 | 497 | 78.1 |
| 7 | Tairajima | 29.69177 | 129.5339 | 2.08 | 243 | 81.9 |
| 8 | Suwanosejima | 29.63833 | 129.7139 | 27.66 | 799 | 60.1 |
| 9 | Akusekijima | 29.465 | 129.5947 | 7.49 | 584 | 81.3 |
| 10 | Kodakarajima | 29.224 | 129.3274 | 1 | 103 | 48 |
| 11 | Takarajima | 29.14444 | 129.2081 | 7.14 | 292 | 32.7 |
| 12 | Yokoatejima | 28.80083 | 128.9947 | 2.76 | 495 | 21.7 |
| 13 | Amamioshima | 28.29611 | 129.3211 | 712.4 | 694 | 65.2 |
| 14 | Kikaijima | 28.3211 | 129.9799 | 56.93 | 214 | 2.9 |

Island numbers correspond to those in Fig. 1.

We first investigated the relationship between species number and island area. A simple linear regression analysis based on log-normalized species number as a dependent variable and log-normalized island area (km^2^) as an explanatory variable was conducted using the lm() function of the *stats* R package.

From the presence/absence datasets of each taxonomic group, pairwise dissimilarity matrices between islands were generated based on three indices, Sørensen, Simpson and nestedness-resultant dissimilarities, using the *betapart* R package [42]. Sørensen dissimilarity evaluates the overall difference between communities based on species composition (β_sør_). The difference between species assemblages, however, results from two distinct situations that the Sørensen dissimilarity index cannot distinguish: (1) differences in assemblages caused by species replacement between communities (spatial turnover: β_sim_) and (2) differences in assemblages caused by differences in species richness (species loss or nestedness-resultant difference: β_nes_). Therefore, β_sim_ and β_nes_ were further estimated using Simpson dissimilarity and nestedness-resultant dissimilarity indices, respectively. Using the *vegan* R package [43], the dissimilarities were visualized in the form of an unrooted dendrogram.

As explanatory variables, seven hypothetical biogeographic boundaries lying between the islands (gaps 1-7 in Fig. 1) were used to examine the significance of the Tokara gap between Akuseki and Kodakara Islands (gap 5 in Fig. 1). Dummy pairwise distance matrices were generated based for each hypothetical boundary in which the distances between islands on the same side of the boundary was given as 0 and the distances between islands across the boundary was given as 1. Pairwise distance matrices of environmental variables, the highest point of island [log_10_(m)], the geographic distance between the highest points of the islands (km), the land area of the island [log_10_(km^2^)], and the proportion of forest (%), were also included in the analyses to explain the correlation between the dissimilarity of species assemblages and environmental factors. These environmental variables were obtained from the National Land Information provided by the Ministry of Land, Infrastructure, Transport and Tourism, Japan. Temperature and precipitation are also important environmental variables that can affect the distribution of organisms and were included in the National Land Information data; however, these were not included in our analyses because these variables were estimated from and correlated with other environmental variables such as the geographic distance or height of the island.

For the regression analyses, the multiple regression on distance matrices (MRM) function implemented in the *ecodist* R package was used [44]. MRM is an extension of the Mantel test and conducts linear regression analysis using distance matrices as dependent and explanatory variables. We first performed simple regression analyses using each of the seven hypothetical biogeographic boundaries or the five environmental variables as the only explanatory variable. We then performed multiple regression analysis using all five environmental variables as explanatory variables. All statistical analyses in this study were conducted in R version 3.3.1 [45].

## RESULTS

### Definitions of the Watase line and the Tokara gap

The Watase line was first proposed by Dr. Yaichiro Okada as a biogeographic boundary between the Palearctic and Oriental realms [13]. This boundary was named after Dr. Shozaburo Watase, who identified a boundary between Tane/Yaku Islands (Is. 1 and 2 in Fig. 1) and Amami Island (Is. 13) based on termite fauna [46]. The biogeographic boundary between Tane/Yaku and Amami Islands was further supported by biogeographic studies on several animal taxa [13,47-49]. The Tokara Archipelago lying between Tane/Yaku and Amami islands was, however, not discussed in these articles. Because limited distribution information for the Tokara Archipelago was available in those days, and limited numbers of species were actually found in the archipelago, many biogeographers did not consider or could not determine which realm each island of the Tokara Archipelago belonged to [13,48,50,51].

We could not find the original article that first proposed the Tokara gap. However, the earliest articles using the term we found were Matsumoto et al. [52] and Kimura [53]. Matsumoto et al. [52] gave a definition of the Tokara gap: a gap between oceanic ridges, the Tane/Yaku Spur and Amami Spur (Fig. 2). Between these spurs, there is a submarine canyon 1000 m below sea level that is deep enough to remain under the sea surface throughout the glacial cycle. However, these spurs and the submarine canyon are distant from the Tokara Archipelago. More importantly, the Tokara gap is not a term for the biogeographical boundary but the name of a bathymetric feature.

### Usage of terms Watase line and Tokara gap

Through a Google Scholar search, we found 108 journal articles in which the terms ‘Watase line’ and/or ‘Tokara gap’ were used. Among them, three contained these terms only in the reference list and were not considered in this study. The number of articles using these terms has increased exponentially (Fig. 3). Among the 105 journal articles, 13 mentioned both the Watase line and the Tokara gap, and 41 and 51 only mentioned the Watase line or the Tokara gap, respectively. All 64 articles that mentioned the Tokara gap were published after the 1990s, and today the Tokara gap is a more frequently used term than the Watase line. Among the 105 articles, 24 put the Watase line or the Tokara gap between Akuseki and Kodakara Islands (gap 5 in Fig. 1) and nine put it another position within the Tokara Archipelago.

**Fig. 3.**
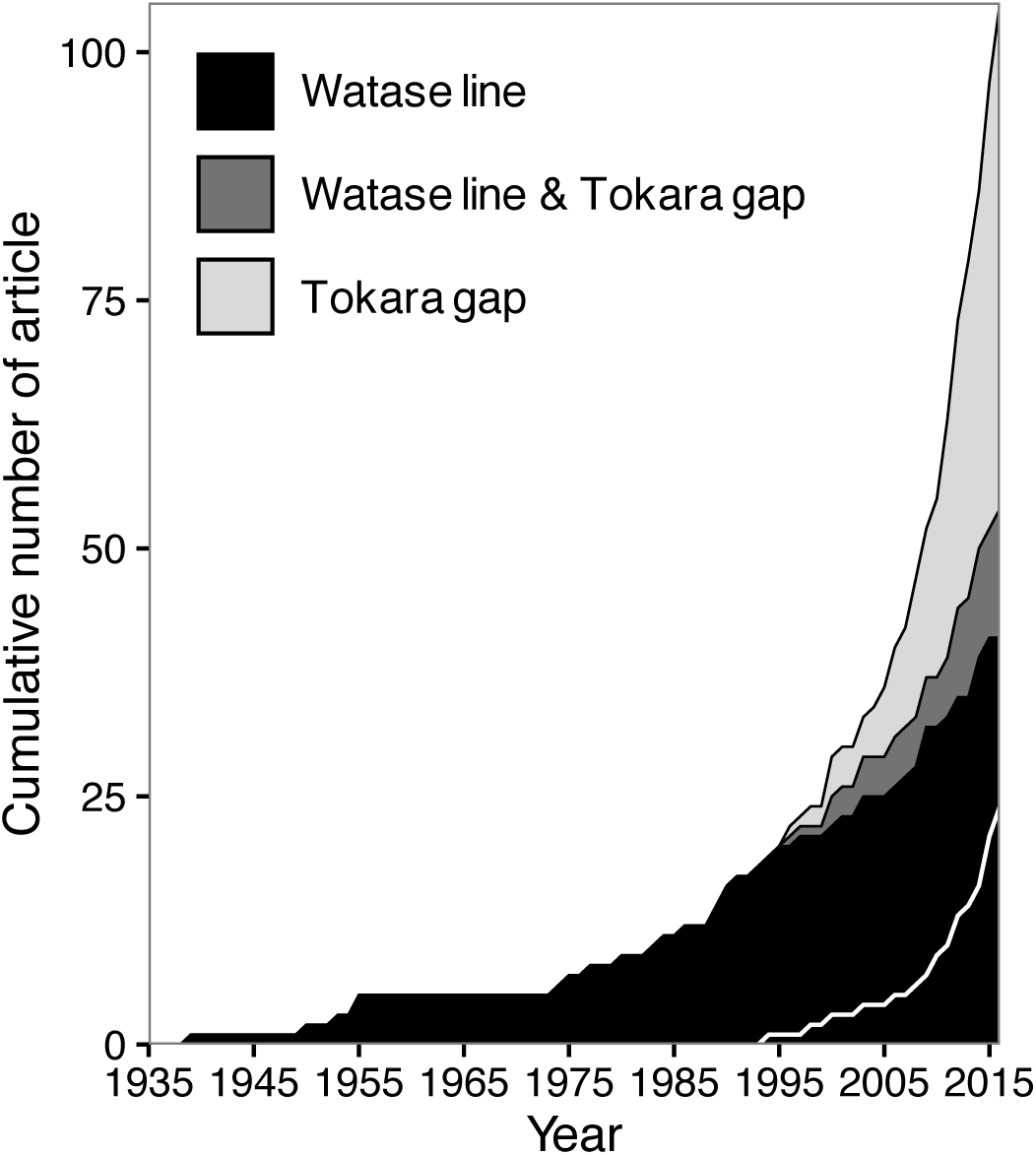
Cumulative number of journal articles in which the terms ‘Watase line’ and/or ‘Tokara gap’ are used. The white line represents the cumulative number of articles that put the biogeographic boundary between Akusekijima and Kodakarajima Islands (gap 5 in Fig. 1).

Among the 105 articles found by the Google Scholar search using ‘Watase line’ and/or ‘Tokara gap’ as keywords, only a single study of flora expressly demonstrated the existence of a biogeographic boundary between Akuseki and Kodakara Islands, while five studies found little genetic differentiation between Akuseki and Kodakara populations of whip scorpions, tree frogs, skinks, cycads and madders [15,16,27,54,55]. The remaining articles performed analyses based on large-meshed sampling skipping the Tokara Archipelago or did not explain the detailed position of the boundary.

### Spatial pattern of species diversity

#### 1. Number of species

The numbers of species in each taxon we collected for analyses were: 125 land snail species from 10 islands (mean species number per island ± sd: 32.3 ± 15.8), 123 ant species from 14 islands (35.4 ± 22.7), 70 butterfly species from 13 islands (28.2 ± 13.7), 69 dragonfly species from 13 islands (19.2 ± 15.6), 17 amphibian species from 10 islands (3.1 ± 3.3), 31 reptile species from 14 islands (6.6 ± 4.5), 53 bird species from 11 islands (25.9 ± 6.4), and 1483 plant species from 14 islands (429.2 ± 285.2) (Fig. 4).

**Fig. 4.**
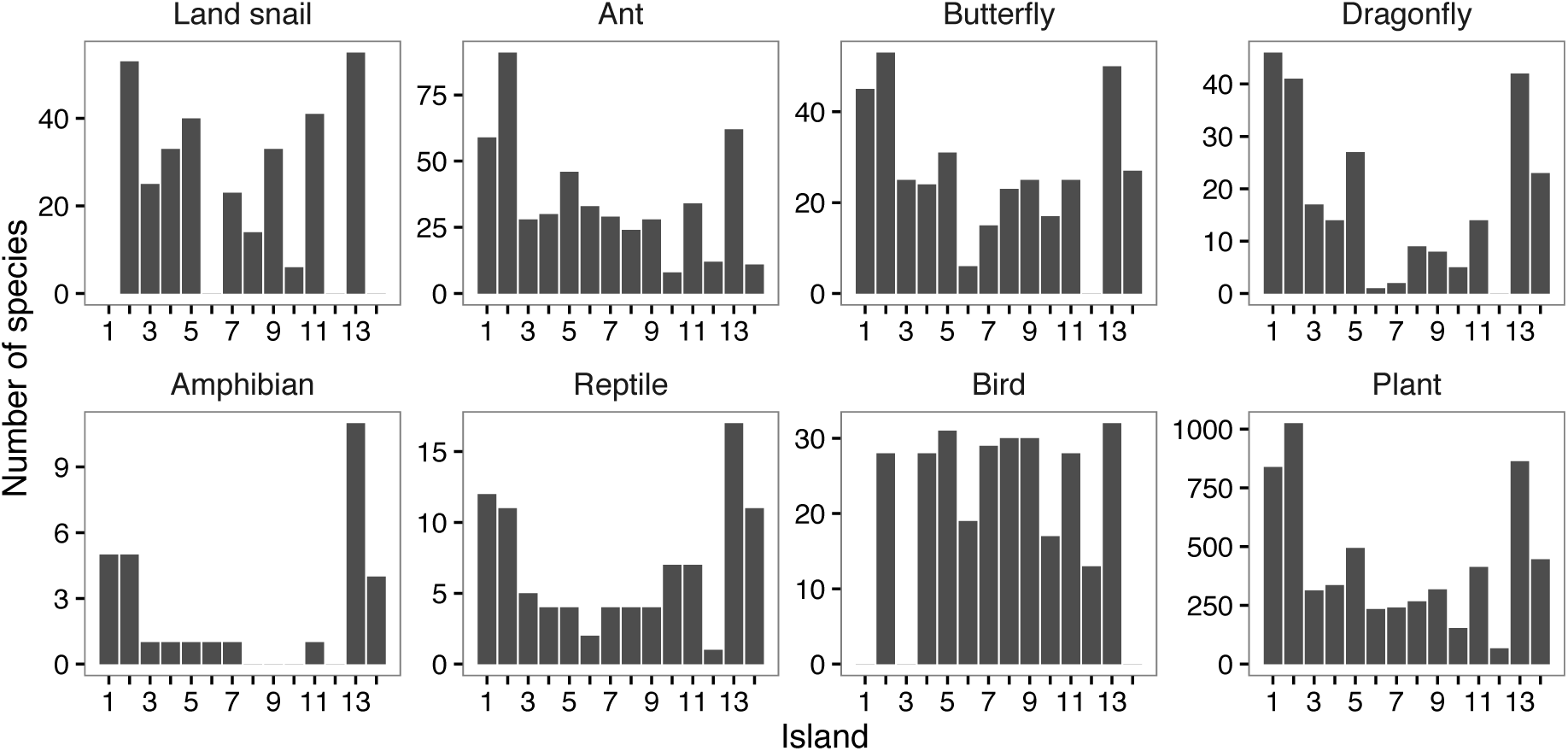
Number of species in each taxon on each island. For both islands for which no distribution data were available and islands on which no species are distributed, the species number is given as 0. Island numbers correspond to those in Table 1 and Fig. 1.

Tane, Yaku and Amami Islands tended to harbour the largest numbers of species except for birds, whose species numbers were almost constant across the islands—around 30 species (Fig. 4). Smaller islands including the Tokara Archipelago and Kuchinoerabujima and Kikaijima Islands, on the other hand, tended to harbour fewer species. A significant correlation between island size and the number of species was found for all taxa except birds (*P* < 0.05, *R*^2^ > 0.4: Fig. 5).

**Fig. 5.**
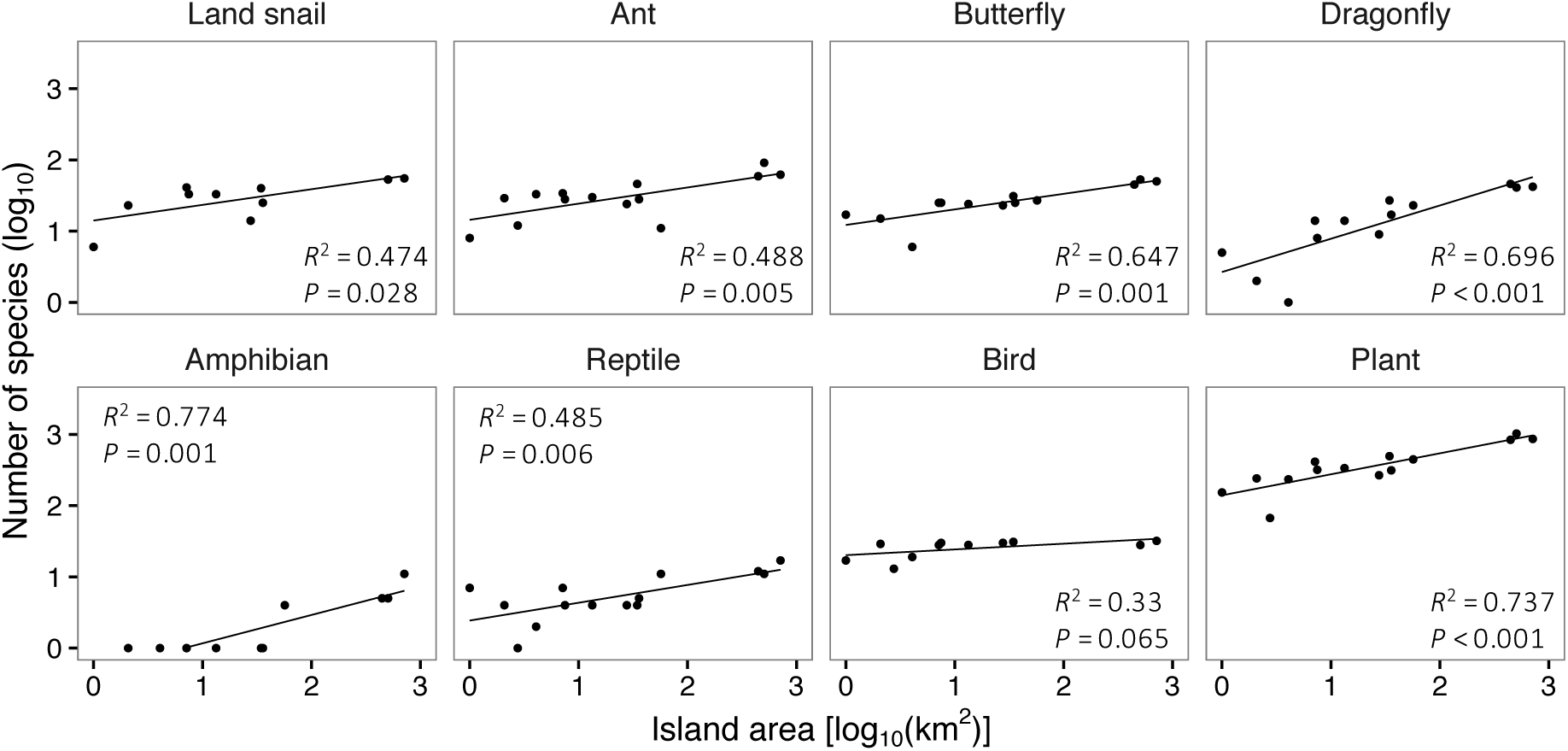
Correlation between island area and species number on the island for each taxon. Regression lines are also presented.

#### 2. Beta diversity

The dissimilarities of species assemblages between islands varied greatly among taxa and the three dissimilarity indices (Additional file 1). These results imply that both spatial-turnover and species-loss significantly contributed to the spatial pattern of species diversity. As expected, for instance, the lower numbers of bird species on Gajajima, Kodakarajima and Yokoatejima Islands (Islands 6, 10 and 12 in Fig. 4) was clearly expressed in the nestedness-resultant dissimilarity (β_nes_) while the Simpson dissimilarity (β_sim_) was seemingly less affected by the number of species (Additional file 1).

In our regression analyses, all seven hypothetical gaps placed in/around the Tokara Archipelago showed significantly positive effects on the dissimilarity of at least a single taxon as follows: gaps 1, 2, 3, 5 and 6 (7) for snails (gaps 6 and 7 are substantially identical because Yokoatejima Is. was ignored), gaps 4, 5, 6 and 7 for ants, gaps 1, 2 and 3 for butterflies, gaps 1, 2 and 6 (7) for amphibians, gaps 1, 2, 3, 4, 5 and 6 for reptiles, gaps 1 and 6 for birds and gaps 4, 5 and 6 for plants (for full information, see Additional file 2). Although many combinations of gaps and beta diversity patterns of taxa showed significant correlations, the determination coefficient (*R*^2^) was relatively small in most cases. For example, gap 5, which is referred to as the Watase line or the Tokara gap, showed an *R*^2^ range of 0.05–0.2, except for reptiles (Table 2 and supplementary table). In reptiles, gap 5 was significantly correlated to β_sør_ with the largest *R*^2^, 0.31. Conversely, gap 1 showed larger *R*^2^ values for amphibian and bird beta diversity patterns. Specifically, the correlations of gap 1 to β_sør_ and β_sim_ for amphibians were *R*^2^ = 0.59 and 1.00, respectively.

**Table 2.** Results of MRM analyses of Sørensen dissimilarity and seven hypothetical gaps for each taxon.

| Model |  | Land snail | Ant | Butterfly | Dragonfly | Amphibian | Reptile | Bird | Plant |
| --- | --- | --- | --- | --- | --- | --- | --- | --- | --- |
| gap 1 | coef | 0.1279 | 0.0503 | 0.0915 | -0.0018 | 0.6181** | 0.2006* | 0.2460 | 0.0419 |
| | $R^2$ | 0.1179 | 0.0214 | 0.0681 | 0.0000 | 0.5945 | 0.1551 | 0.2838 | 0.0179 |
| gap 2 | coef | 0.0752 | 0.0204 | 0.0714 | -0.0145 | 0.2986* | 0.1365* | 0.0661 | 0.0138 |
| | $R^2$ | 0.0442 | 0.0038 | 0.0436 | 0.0011 | 0.1388 | 0.0765 | 0.0303 | 0.0021 |
| gap 3 | coef | 0.0595 | 0.0326 | 0.0662* | 0.0353 | 0.0188 | 0.1244* | -0.0078 | 0.0163 |
| | $R^2$ | 0.0277 | 0.0097 | 0.0374 | 0.0063 | 0.0006 | 0.0643 | 0.0005 | 0.0029 |
| gap 4 | coef | 0.0518 | 0.0698* | -0.0105 | -0.0124 | 0.0585 | 0.2435*** | 0.0021 | 0.0391 |
| | $R^2$ | 0.0210 | 0.0444 | 0.0009 | 0.0008 | 0.0053 | 0.2465 | 0.0000 | 0.0168 |
| gap 5 | coef | 0.1615* | 0.1123** | -0.0101 | -0.0348 | 0.0585 | 0.2716*** | 0.0618 | 0.0811* |
| | $R^2$ | 0.2043 | 0.1154 | 0.0009 | 0.0062 | 0.0053 | 0.3073 | 0.0301 | 0.0725 |
| gap 6 | coef | 0.2000* | 0.1702* | 0.0232 | -0.0328 | 0.2367 | 0.1266* | 0.1968* | 0.1217* |
| | $R^2$ | 0.2014 | 0.2450 | 0.0037 | 0.0045 | 0.0803 | 0.0618 | 0.2687 | 0.1509 |
| gap 7 | coef | 0.2000* | 0.1581* | 0.0232 | -0.0328 | 0.2367 | 0.1399 | 0.2048 | 0.0361 |
| | $R^2$ | 0.2014 | 0.1776 | 0.0037 | 0.0045 | 0.0803 | 0.0633 | 0.1966 | 0.0111 |

For the results using other dissimilarity indices, see Additional file 2. coef: coefficient. (\* $P <$
0.05, \*\* $P < 0.01$ , \*\*\* $P < 0.001$ )

#### 4. Environmental factors for the spatial pattern of species diversity

All four environmental variables used in this study showed significant effects on the dissimilarity of species assemblies for all taxa and dissimilarity indices (Table 3 and Additional file 2). Here, we mainly mention the results of analyses based on β_sør_ (for full results, see Additional file 2). In land snails, amphibians, birds and plants, the area and geographic distance of the islands showed significant positive correlations; in ants and reptiles, the area, geographic distance and forest cover of the islands showed significant positive correlations; in butterflies and dragonflies, the area of the islands showed a significant positive correlation (Table 3). Throughout the taxa, the dissimilarity of island area was positively correlated with overall species dissimilarity (β_sør_).

**Table 3.** Results of MRM analyses of Sørensen dissimilarity and environmental variables for each taxon.

| Model | Land snail | Ant | Butterfly | Dragonfly | Amphibian | Reptile | Bird | Plant |
| --- | --- | --- | --- | --- | --- | --- | --- | --- |
| Area coef | 0.1499** | 0.0909** | 0.1068** | 0.1380** | 0.2483* | 0.0967* | 0.1547** | 0.1216*** |
| $R^2$ | 0.3803 | 0.1590 | 0.2020 | 0.2008 | 0.1822 | 0.0820 | 0.4447 | 0.3426 |
| Height coef | 0.2525 | 0.2044 | 0.0220 | 0.0333 | 0.0329 | 0.1303 | 0.1337 | 0.0938 |
| $R^2$ | 0.1810 | 0.1083 | 0.0012 | 0.0017 | 0.0004 | 0.0200 | 0.0450 | 0.0274 |
| Forest coef | 0.0037 | 0.0033* | -0.0006 | -0.0017 | 0.0007 | 0.0030* | 0.0020 | 0.0016 |
| $R^2$ | 0.1005 | 0.2026 | 0.0072 | 0.0308 | 0.0019 | 0.0773 | 0.0448 | 0.0596 |
| Geodis coef | 0.0020*** | 0.0010* | 0.0006 | -0.0001 | 0.0031** | 0.0021*** | 0.0020** | 0.0008* |
| $R^2$ | 0.4400 | 0.2009 | 0.0613 | 0.0009 | 0.3569 | 0.3902 | 0.4241 | 0.1338 |
| Area coef | 0.0830* | 0.0742** | 0.0978** | 0.1620*** | 0.1770* | 0.0228 | 0.1264** | 0.1262*** |
| Height coef | 0.0987 | 0.1142 | -0.0377 | -0.0753 | 0.1497 | 0.1083 | -0.0568 | -0.0220 |
| Forest coef | 0.0017 | 0.0028* | -0.0005 | -0.0002 | -0.0029 | 0.0005 | 0.0009 | 0.0019 |
| Geodis coef | 0.0014** | 0.0005 | 0.0004 | -0.0006 | 0.0033*** | 0.0020*** | 0.0012* | 0.0001 |
| $R^2$ | 0.6491 | 0.4501 | 0.2259 | 0.2410 | 0.5034 | 0.4143 | 0.6138 | 0.4410 |

For the results using other dissimilarity indices, see Additional file 2. Geodis: geographic
distance, coef: coefficient. (\* $P < 0.05$ , \*\* $P < 0.01$ , \*\*\* $P < 0.001$ )

However, the determination coefficients (*R*^2^) for each environmental variable varied among taxa. In land snails, the geographic distance of the islands was significantly correlated with β_sør_ with the highest *R*^2^ (0.44). The regression analysis based on β_sim_ and geographic distance showed a larger *R*^2^ (0.71). In ants, all simple regression analyses showed relatively small *R*^2^ values. The multiple regression analysis based on β_sør_ and all four environmental variables showed a larger *R*^2^ of 0.45. Additionally, in butterflies and dragonflies, all regression analyses showed small *R*^2^ values. Even multiple regression analyses based on all four variables showed *R*^2^ values below 0.25. In amphibians, geographic distance showed the highest *R*^2^ (0.36), but this was smaller than the *R*^2^ of hypothetical gap 1 (0.59; Table 2). Similarly, in reptiles, geographic distance showed the highest *R*^2^ (0.39). In birds, the area and geographic distance of the islands showed the largest *R*^2^ values (0.44 and 0.42, respectively). In plants, the area of the islands showed the largest *R*^2^ (0.34).

The signs (positive/negative) of the correlation coefficients between island area and β_sim_ varied among taxa, even those showing *P* values lower than 0.05 (Additional file 2). However, these analyses tended to show smaller *R*^2^ values and/or correlation coefficients of almost zero, indicating they had no significant biogeographical implications.

## DISCUSSION

### ‘Watase line’ and ‘Tokara gap’ as terms

Our documentary search revealed that the Watase line was proposed as a biogeographic boundary between Tane/Yaku and Amami Islands. On the other hand, the Tokara gap is the name of a bathymetric feature, a deep submarine canyon between Tane/Yaku and Amami Islands (Fig. 2). It is highly possible that the bathymetric feature that is the Tokara gap is responsible for the biogeographic boundary of the Watase line; a sea barrier formed by the deep submarine canyon between the Tane/Yaku and Amami Spurs (the Tokara gap) inhibited the dispersal of terrestrial organisms for a long period, which led to the differentiation of fauna and flora between Tane/Yaku and Amami Islands (the Watase line).

Today, the Watase line (Tokara gap) is generally put between Akuseki and Kodakara Islands of the Tokara Archipelago, however, it seems incorrect because the Watase line was put between Tane/Yaku and Amami Islands and the Tokara Archipelago was not considered in the original article, and furthermore, the Tokara gap—a deep submarine canyon—does not lie between Akuseki and Kodakara Islands (Fig. 2). Through the document search in this study, we found three possible causes for the misplacement of the boundary; (1) the position of a sea strait, (2) the formation of a land bridge and (3) the distribution of pit vipers.

(1) In several biogeographic studies, it was noted that the Tokara strait (Tokara tectonic strait) lies between Akuseki and Kodakara Islands, and has acted as a geographical barrier to terrestrial organisms [16,20,56-60]. These studies declared that the barrier, the Tokara strait, has existed since the Pliocene, and referred to it as the Tokara gap (gap 5 in Fig. 1). However, as mentioned above, the Tokara gap, the deep submarine canyon (-1000 m in depth), does not lie between Islands of the Tokara Archipelago. Furthermore, the position of the Tokara strait is not strictly defined but varies depending on the context [61].

(2) A land connection between Amami and Kodakara Islands has been depicted in the figures of several articles (e.g. Nakamura et al. 2012; Kumekawa et al. 2014). If the islands had any land connections, the terrestrial and freshwater biotas of Kodakara Island would share more species with Amami Island than with Akuseki Island. This paleogeographic inference should support the idea of a biogeographic boundary between Kodakara and Akuseki Islands. However, no evidence for the land bridge hypothesis was mentioned in their arguments. To our knowledge, the only geographic factor that implies a land bridge connection between Amami and Kodakara Islands is the distribution of Ryukyu limestone. This is a reef-building limestone deposited during the Pleistocene, reflecting the expanse of shallow sea during the period. According to Kizaki [61] and Kato [63], this limestone is continuously distributed between Amami and Kodakara Islands, and was possibly deposited along a land bridge once formed between the islands. However, the distribution data for Ryukyu limestone have always been referred to as “unpublished data”, and we could not find any published articles that report the details of the data. Therefore, the land bridge hypothesis is unevaluable unless a study on the limestone distribution is published.

(3) The most symbolic and frequently referred to taxon that represents the existence of a biogeographic boundary is *Protobothrops*, a genus of venomous pit vipers in the Ryukyu Archipelago [64-66]. These pit vipers are widely distributed throughout the Ryukyu Archipelago, and Kodakara Island is the northernmost island on which a *Protobothrops* species is found [33]. Based on the vipers’ distribution, Hikida et al. [65] suggested the existence of a biogeographic boundary between Kodakara and Akuseki Islands. However, the distribution pattern varies among taxa even within snake species [33]. Although the distribution pattern of vipers suggests the existence of a geological or ecological barrier for vipers between Kodakara and Akuseki Islands, the idea cannot be applied to other organisms that have different ecological characters.

In addition, despite the growing number of articles that depict the boundary lying between Akuseki and Kodakara Islands (Fig. 3), few of them have investigated the biogeography using samples or data collected from the Tokara Archipelago. It means that, in most studies, the location of the biogeographic boundary was not important or it was just taken from other articles without verification. This could have enhanced the spread of the idea that the Tokara gap lies between Akuseki and Kodakara Islands.

### Biogeography in the Tokara Archipelago

The regression analyses of species number and area of the islands showed clear positive correlation between them in all taxa except for bird (Fig. 5). This finding that larger islands harbour more species fits one of the general laws of island biogeography [2,3].

In our beta diversity analyses, all seven hypothetical gaps placed in/around the Tokara Archipelago showed significantly positive effects on the dissimilarity of at least a single taxon (Table 2 and Additional file 2). Although, no gaps showed significant effects across all eight taxa. These results suggest that there is no prominent biogeographic boundary around the Tokara Archipelago, but that the biota changes gradually on a spatial scale. The Watase line or the Tokara gap misplaced between Akuseki and Kodakara Islands does not represent the biogeographic patterns of fauna and flora in this region.

The beta diversity pattern of amphibians was largely shaped by the distribution of *B. japonica*; thus, gap 1, which corresponds to the northern limit of the distribution, should well explain the beta diversity pattern of amphibians. Conversely, except for ants and amphibians, a single environmental variable could explain the beta diversity pattern better than any hypothetical gap considered in this study, showing larger determination coefficients (*R*^2^) (Table 3 and Additional file 2). In addition, multiple regression analysis applying all four environmental variables showed an *R*^2^ larger than that of any hypothetical gap in ants. In particular, it is obvious that the areas and geographic distances of the islands are determining factors for the beta diversity patterns of the fauna and flora in this region (Table 3), suggesting that the spatial pattern of species diversity in this region obeys the principles of island biogeography, distance decay and the species-area relationship, rather than the misplaced historical biogeographic boundary, the Tokara gap.

### Effects of preconception on the biogeographic debates

Several biogeographic studies have performed beta diversity analyses based on the fauna and flora of the Ryukyus including the Tokara Archipelago, and demonstrated the presence of a major biogeographic boundary between Akuseki and Kodakara Islands. However, our analyses of beta diversity in the Tokara Archipelago did not support this idea. Here, we compare and discuss the discordance between present and previous studies.

Ichikawa et al. [26] argued that the Watase line (gap 5), which was put between Akuseki and Kodakara Islands, had a significant effect on snail diversity. However, as clarified above, this is not the sole boundary that shapes the spatial pattern of species diversity for snails. Hirao et al. [32] analysed the association of the spatial pattern of butterfly fauna to the Tokara gap (gap 5) and the Kerama gap—which is placed in the southern Ryukyus. They showed that these gaps had a significant effect on the butterfly fauna, especially on the nestedness dissimilarity. In our analysis, in contrast, a significant effect was not found for gap 5. This could be due to differences in analysis: we only focused on the islands in/around the Tokara Archipelago and the Tokara gap whereas Hirao et al. [32] studied the entire region of the Ryukyu Archipelago and simultaneously analysed the effects of both the Tokara and Kerama gaps. In the analyses based on reptile distribution, six hypothetical gaps showed significant effects. Eleven reptile species are distributed on nine islands of the Tokara Archipelago, and six islands represent the northern/southernmost populations of eight species [33,36]. Therefore, almost every hypothetical gap (gaps 1-6) corresponds to the distribution boundary of a certain reptile species. As stated above, the pit viper genus *Protobothrops*, the northern distribution limit of which is Kodakara Island, is a key genus supporting the idea of a biogeographic boundary between Akuseki and Kodakara Islands [64-66]; however, another species shows a different position as the distribution limit. Nakamura et al. [20] and Kubota et al. [25] investigated the correlation between the Tokara gap (gap 5) and the spatial pattern of flora in the Ryukyu Archipelago, and demonstrated that the gap had a significant effect on the flora pattern. They suggested that the large floristic difference between Akuseki and Kodakara Islands implies the existence of a historical barrier, the Tokara gap. However, again, this is not the sole boundary; three hypothetical boundaries analysed in this study showed significant contributions to the floristic differentiation among islands of the Tokara Archipelago.

It is noteworthy that all hypothetical boundaries examined in our analyses had a significant effect on the beta diversity pattern, while the abovementioned studies focused on just one of them, between Akuseki and Kodakara Islands. Thus, it is highly possible that their arguments were strongly biased by the preconception that the boundary lay between Akuseki and Kodakara Islands.

## CONCLUSIONS

Neither our document search nor our biogeographic analyses supported the presence of a clear biogeographic boundary between Akuseki and Kodakara Islands. Our biogeographic analyses suggested that the biota varies among islands, and a sea strait between Akuseki and Kodakara Islands could only partially explain the beta diversity pattern of this region. In other words, the Tokara Archipelago cannot be simply dichotomized into Palearctic and Oriental realms. The widespread idea of a biogeographic boundary (the Watase line or the Tokara gap) between Akuseki and Kodakara Islands is baseless, and we discourage biogeographic reconstruction relying on this misconception. Furthermore, the islands of the Tokara Archipelago are thought to be oceanic islands that never had land-bridge connections to other islands because they are volcanic in origin and developed from the deep sea floor [66,67]. In this case, the biota in the Tokara Archipelago should have never been affected by a geohistory of land-bridge formation and submergence, but consists of species that achieved dispersal over the sea. At present, it is adequate to put the boundary between Tane/Yaku and Amami Islands, and the Tokara Archipelago seems to be a gap between the Palearctic and Oriental realms.

Besides demonstrating the necessity for revision of the biogeography in the Tokara Archipelago, this study demonstrates the pitfalls and risks of preconception in biogeographic debate. Specifically, it reveals a vicious cycle: preconception affects the design or interpretation of biogeographic analyses and subsequent biased results further perpetuate this preconception.

**Fig. 6.**
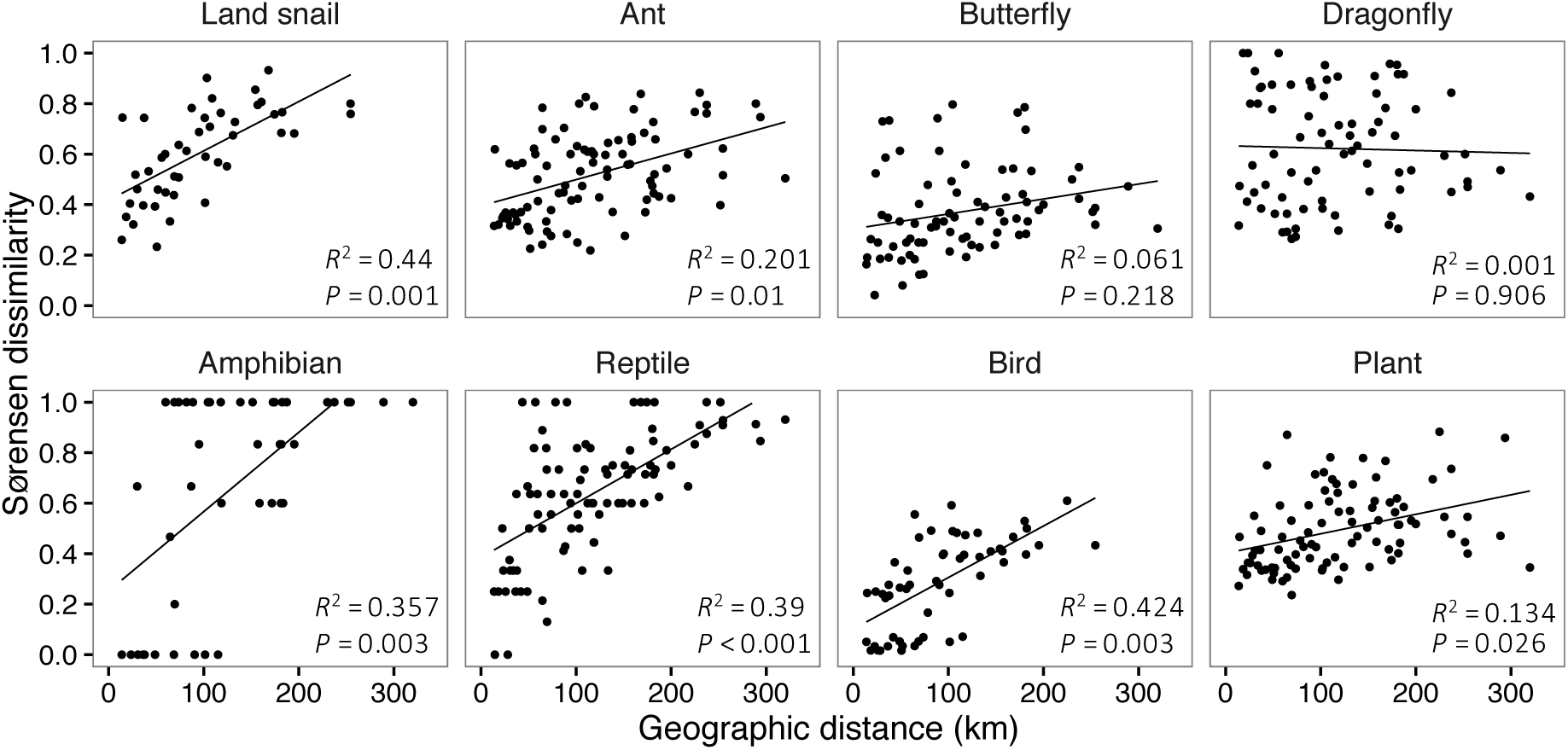
Correlation between geographic distance and Sørensen dissimilarity for each taxon. Regression lines are also presented.

**Fig. 7.**
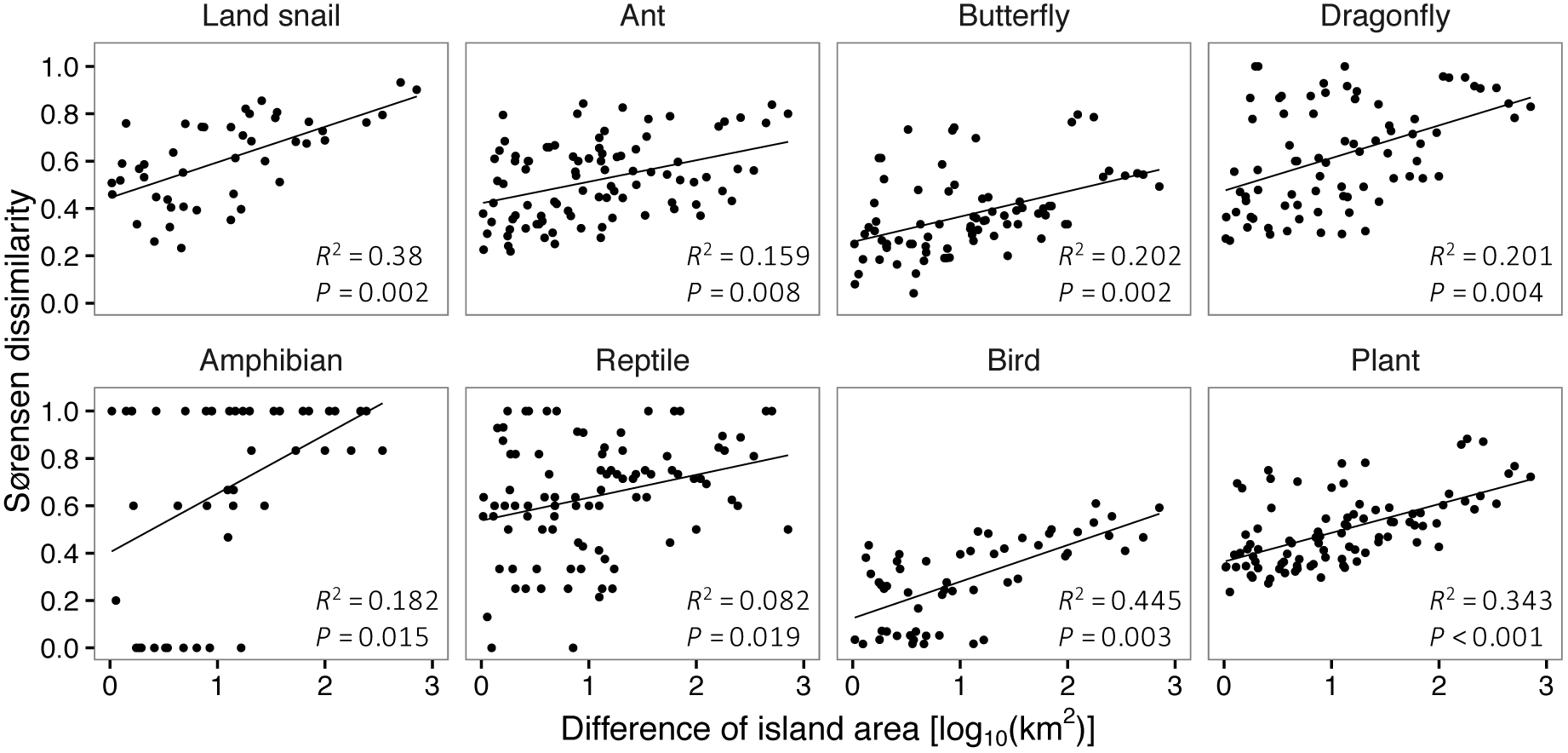
Correlation between the difference of island area and Sørensen dissimilarity for each taxon. Regression lines are also presented.

## DECLARATIONS

### Ethics approval and consent to participate

Not applicable.

### Consent for publication

Not applicable.

### Availability of data and material

The datasets used in this study are available from articles or public database referred in this manuscript.

### Authors’ contributions

S.K. conceived the study; S.K. and T.I. collected documents; S.K. compiled and analysed data; S.K. led the writing and T.I. reviewed the manuscript.

## Acknowledgements

The authors are grateful to Prof. Koji Tojo (Shinshu University), Dr. Yuh Shiwa (Iwate Medical University) and Dr. Ryohei Furukawa (Iwate Medical University) for their assistance with data collection and analyses.

## ADDITIONAL FILES

### Additional file 1

Dendrograms representing dissimilarity of species composition of land snail, ant, butterfly, dragonfly, amphibian, reptile, bird and plant on islands in/around the Tokara Archipelago. Sørensen, Simpson and nestedness-resultant dissimilarity indices were applied to estimate the dissimilarity. Numbers on each dendrogram represent the island number listed in Table 1 of this article. (file name: Additionalfile1.docx, file size: 465 kb)

### Additional file 2

Results of simple and multiple regression analyses based on hypothetical gaps and environmental factors. (file name: Additionalfile2.xls, file size: 74 kb)

